## Supplementary figures and images for "Abundant genetic variation is retained in many laboratory schistosome populations"

### Supplemental Figure 1

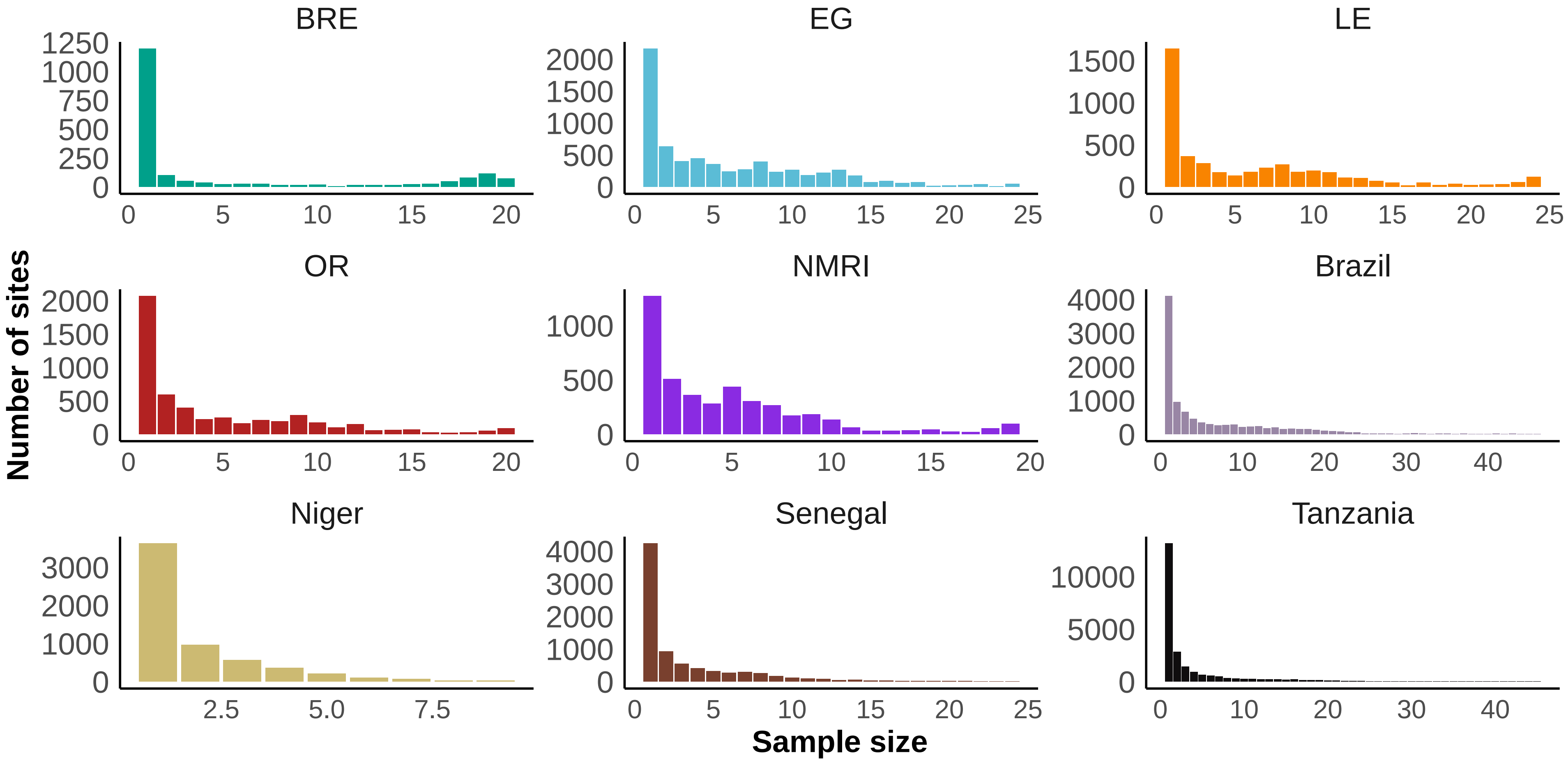

### Supplemental Figure 2

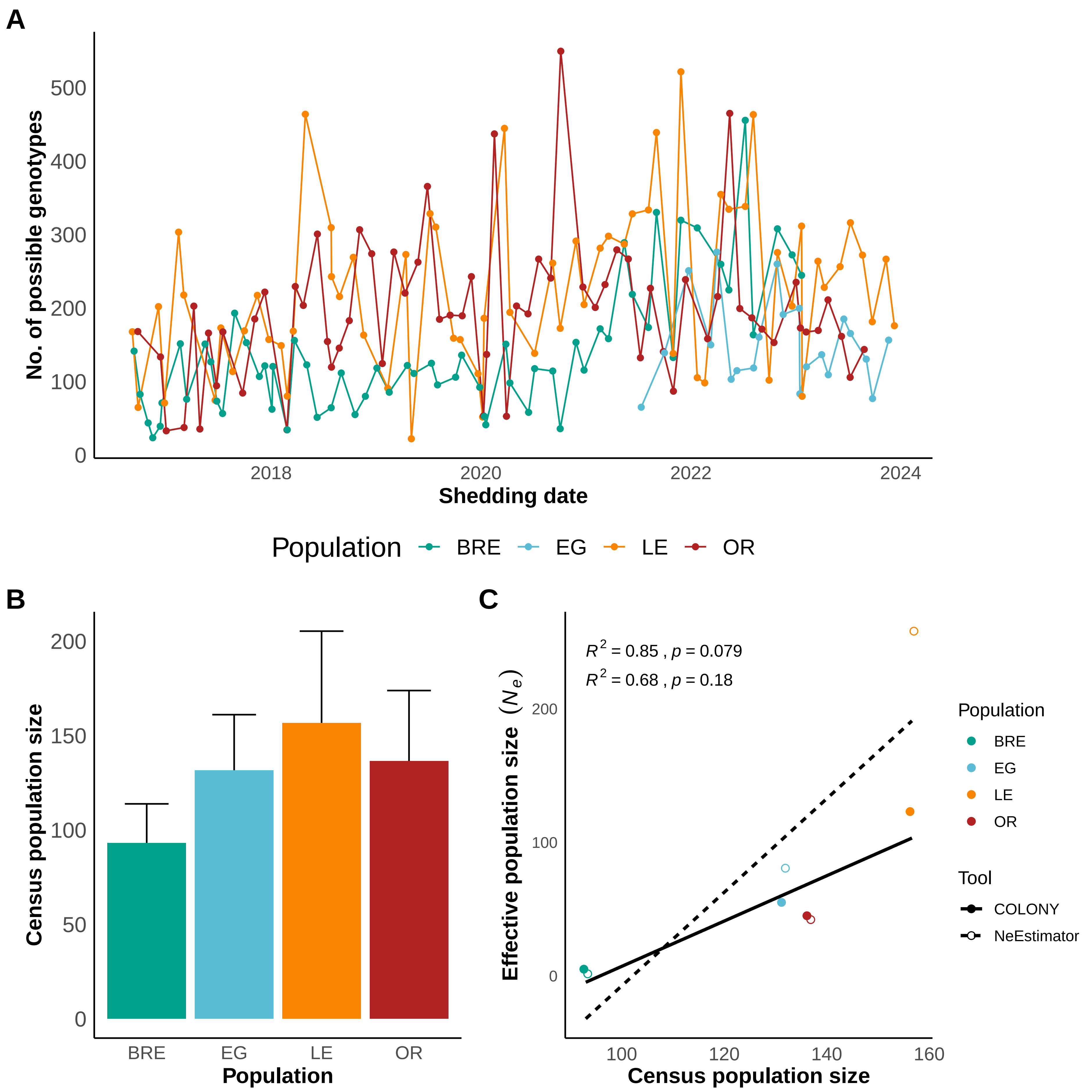

### Supplemental Figure 3

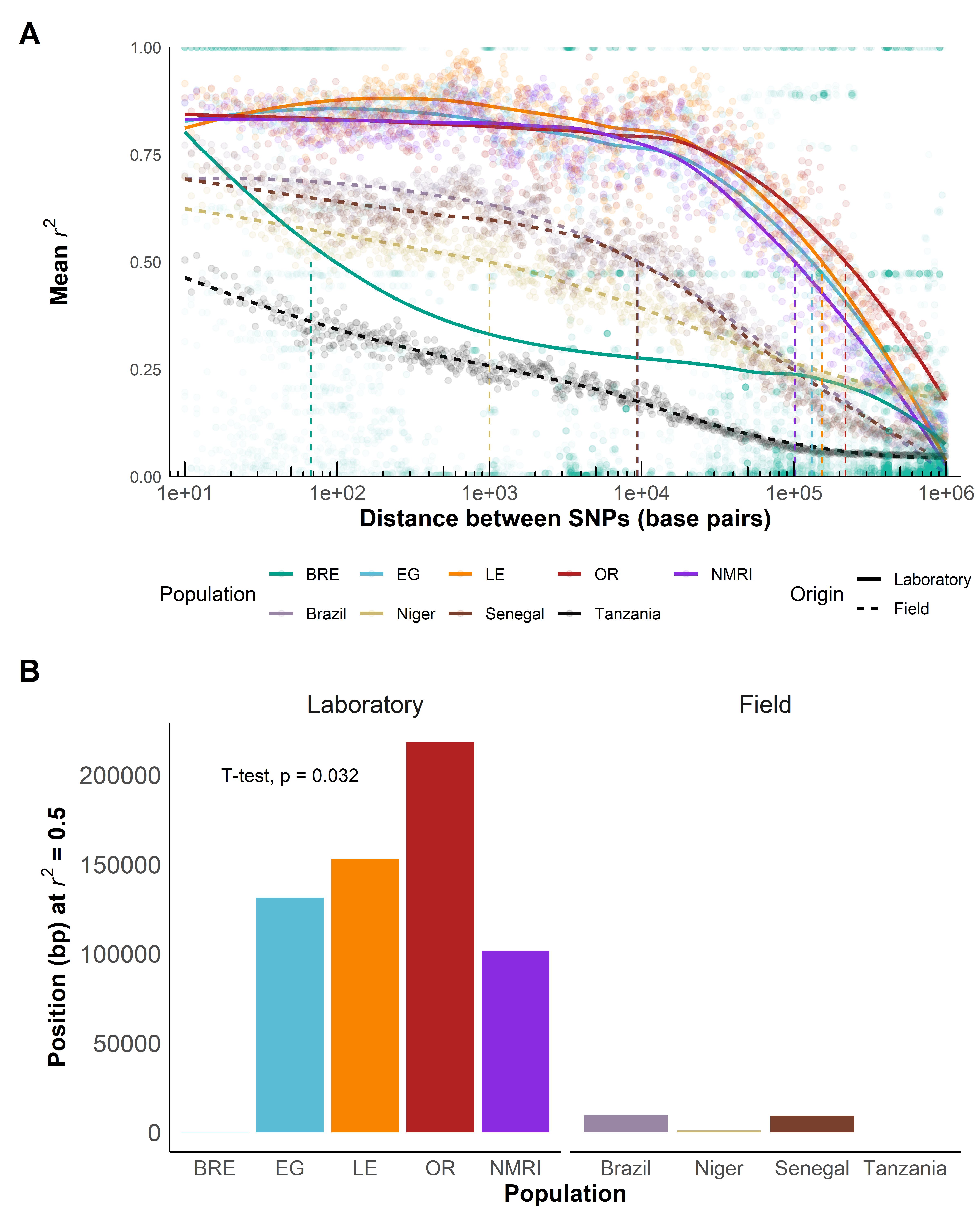
